# Cell-type specific contributions to theta-gamma coupled rhythms in the hippocampus

**DOI:** 10.1101/2024.03.05.583493

**Authors:** Spandan Sengupta, Afroditi Talidou, Jeremie Lefebvre, Frances K Skinner

## Abstract

Distinct inhibitory cell types participate in cognitively relevant nested brain rhythms, and particular changes in such rhythms are known to occur in disease states. Specifically, the co-expression of theta and gamma rhythms in the hippocampus is believed to represent a general coding scheme, but cellular-based generation mechanisms for these coupled rhythms are currently unclear. We develop a population rate model of the CA1 hippocampus that encompasses circuits of three inhibitory cell types (bistratified cells, parvalbumin (PV)-expressing and cholecystokinin (CCK)-expressing basket cells) and pyramidal cells to examine this. We constrain parameters and perform numerical and theoretical analyses. The theory, in combination with the numerical explorations, predicts circuit motifs and specific cell-type mechanisms that are essential for the co-existence of theta and gamma oscillations. We find that CCK-expressing basket cells initiate the coupled rhythms and regularize theta, and PV-expressing basket cells enhance both theta and gamma rhythms. Pyramidal and bistratified cells govern the generation of theta rhythms, and PV-expressing basket and pyramidal cells play dominant roles in controlling theta frequencies. Our circuit motifs for theta-gamma coupled rhythm generation could be applicable to other brain regions.

**AUTHOR SUMMARY:** There are many different types of inhibitory cells in our brains that are differentially affected in disease. Concomitantly, coupled rhythms change in particular ways with disease. To help understand cell-type specific changes in coupled rhythms, we develop a mathematical network model that is both respective of the cell type and also amenable to analyses. We focus on theta-gamma coupled rhythms in the hippocampus and include three different inhibitory cell types in our model circuits. By combining theoretical analysis and numerical explorations, we find distinct contributions of these inhibitory cell types to coupled rhythms, and predict motifs that are essential for the expression of theta-gamma coupled rhythms. Moving forward, we can leverage our model insights to help unravel cell-type contributions in disease states.

## INTRODUCTION

Oscillatory activities of wide-ranging frequencies are ubiquitous in many brain structures (Buzsáki, 2006). Theta oscillations, ranging from 3 to 12 Hz, are prominent local field potential (LFP) rhythms that are most robustly recorded from the CA1 region (Buzsáki, 2002). This rhythm plays an essential role in memory and spatial navigation (Buzsáki & Moser, 2013), with its frequency being related to the animal’s kinematics (Kropff, Carmichael, Moser, & Moser, 2021). High (7-12 Hz) and low (4-7 Hz) ranges of theta frequencies dependent on atropine sensitivity have long been observed (Buzsáki, 2002; Kramis, Vanderwolf, & Bland, 1975), and can be distinctly elicited by social or fearful stimuli, respectively (Tendler & Wagner, 2015). A single theta cycle has been considered to be ‘a functional unit capable of representing distinct temporal-spatial content at different phases’ (M. A. Wilson, Varela, & Remondes, 2015), and higher frequency gamma (≈20-100 Hz) oscillations are nested within these theta cycles (e.g., see Scheffer-Teixeira et al. (2012)). It has been suggested that such theta-gamma coupling is a general coding scheme with information processing implications (Colgin, 2015; Jensen & Colgin, 2007; Lisman, 2005). Dynamic modulation of theta-gamma coupled rhythms during sleep (Bandarabadi et al., 2019), visual exploration (Kragel et al., 2020) and association with working memory exists, and it is thus not surprising that there is increasing evidence of specific changes in theta-gamma coupling with memory impairments, disease and its progression (Goutagny et al., 2013; Hamm, Héraud, Cassel, Mathis, & Goutagny, 2015; Karlsson, Lindenberger, & Sander, 2022; Kitchigina, 2018; Musaeus, Nielsen, Musaeus, & Høgh, 2020; Zhang et al., 2016). Yet, it remains unclear what cell circuit motifs and mechanism(s) are responsible for these co-expressed rhythms.

The plethora of various interneurons or inhibitory cell types with their particular biophysical characteristics and connectivities in the hippocampus make it challenging to figure out their particular contributions to functionally relevant theta and gamma activities (Buzsáki, 2006; Fishell & Kepecs, 2020; Freund & Buzsáki, 1996; Kepecs & Fishell, 2014; Klausberger & Somogyi, 2008; McBain & Fisahn, 2001; Pelkey et al., 2017). However, as theta-gamma coupled rhythms reflect cognitive processing, are potential disease biomarkers, *and* involve various interneuron types, we cannot ignore this inhibitory diversity. For example, in trying to understand the contributions of specific inhibitory cell types, it has long been noted that there is a separation between perisomatically and dendritically targeting interneuron types onto pyramidal cells (Freund & Buzsáki, 1996; McBain & Fisahn, 2001) leading to consideration of a dichotomy in inhibitory control of pyramidal cell excitability. Further, Freund (2003) has described another dichotomy of perisomatically targeting cholecystokinin (CCK)-expressing and parvalbumin (PV)-expressing basket cells in a ‘rhythm and mood’ fashion (Freund, 2003; Freund & Katona, 2007) – PV-expressing basket cells contribute in a precise clockwork fashion, and CCK-expressing basket cells in a highly modulatory way. Using and combining modern technologies, the distinctness of interneurons is being unravelled (Gouwens et al., 2020; Harris et al., 2018; Kessaris & Denaxa, 2023; Ratliff & Batista-Brito, 2020; Yao et al., 2021). Twenty-eight well-defined interneuron types have been identified and bistratified cells, PV-expressing basket cells and CCK-expressing basket cells are distinguishable. These collective studies help us bridge cell-type focused experiments with theoretical and modeling endeavours.

Many mathematical models have been used to help determine mechanisms underlying theta, gamma and theta-gamma coupled rhythms in the hippocampus (Ferguson & Skinner, 2015). For gamma rhythms, ING (interneuron network gamma) and PING (pyramidal interneuron network gamma) mechanisms have long been proposed (Whittington, Traub, Kopell, Ermentrout, & Buhl, 2000). For theta and theta-gamma coupled rhythms, network models have mostly focused on the dynamics between oriens-lacunosum-moleculare (OLM) interneurons, fast-spiking interneurons, and pyramidal cells (Ferguson & Skinner, 2015). Recently, building from tight experimental linkages, we showed that a large enough network of pyramidal cells could initiate theta rhythms (Chatzikalymniou, Gumus, & Skinner, 2021). However, in general, high dimensional spiking network model systems challenge our capacity to extract explicit mechanisms. To circumvent this, we here build and analyze a cell-type specific mean field model that combines pyramidal cells and three distinct inhibitory cell types (bistratified cells, PV-and CCK-expressing basket cells) that were found to be essential for theta-gamma coupled rhythms from cell-type and cell-type interaction perspectives (Bezaire, Raikov, Burk, Vyas, & Soltesz, 2016; Chatzikalymniou, Sengupta, Lefebvre, & Skinner, 2022). This model leverages population-scale description of neural activity with experimentally constrained parameters to gather insight about theta-gamma coupling and theta frequency control in the hippocampus, while providing testable predictions.

We perform high-throughput simulations and obtain multiple constrained parameter sets for theta-gamma coupled rhythms indicating degeneracy. We systematically analyze the contribution of individual cell types involved in theta-gamma coupled oscillations by quantifying changes in spectral characteristics resulting from targeted stimulation of individual cell types, and we also examine whether particular cell types play dominant roles in the control of theta frequency. We perform theoretical analysis which, in combination with numerical explorations, predicts cell types and motifs essential for co-existence of theta and gamma oscillations. We find that circuit motifs of pyramidal and bistratified cells play a governing role in the theta rhythms, with CCK-expressing and PV-expressing basket cells together with pyramidal cells contributing to gamma rhythms. Taken together, our results show that CCK-expressing basket cells act to initiate theta-gamma coupled rhythms by their ability to control pyramidal cell activity via disinhibition, and we present a general two-phase process by which theta-gamma coupled rhythms arise in network motifs.

## RESULTS

### Building and exploring a population rate model (PRM) of theta-gamma coupled rhythms

In order to disambiguate the respective role of cell types and their mutual connectivity motifs in the generation and control of theta-gamma coupled rhythms in the CA1 hippocampus, we developed a reduced, system-level mean field population rate model (PRM). This model combines connections and cell types that were identified as critical for the existence of theta-gamma rhythms based on previous work using more detailed circuit models with nine different cell types – pyramidal cells (**PYR**), bistratified cells (**BiC**), PV-expressing basket cells (**PV**), CCK-expressing basket cells (**CCK**), axo-axonic cells, Schaeffer collateral-associated cells, ivy cells, neurogliaform cells and OLM cells. (Bezaire et al., 2016; Chatzikalymniou et al., 2021, 2022). Specifically, we previously showed that theta rhythms are lost with the removal of certain connections, and we rationalized the use of a four cell-type (PYR, BiC, CCK, PV) microcircuit (Bezaire et al., 2016; Chatzikalymniou et al., 2022) to examine theta-gamma coupled rhythms in CA1 hippocampus. In FIGURE 1 **A-B**, we show schematics of the cell types and interconnections used in previous detailed microcircuit models of the CA1 hippocampus (**A**), and the four cell-type microcircuit used here (**B**).

**Figure 1.**
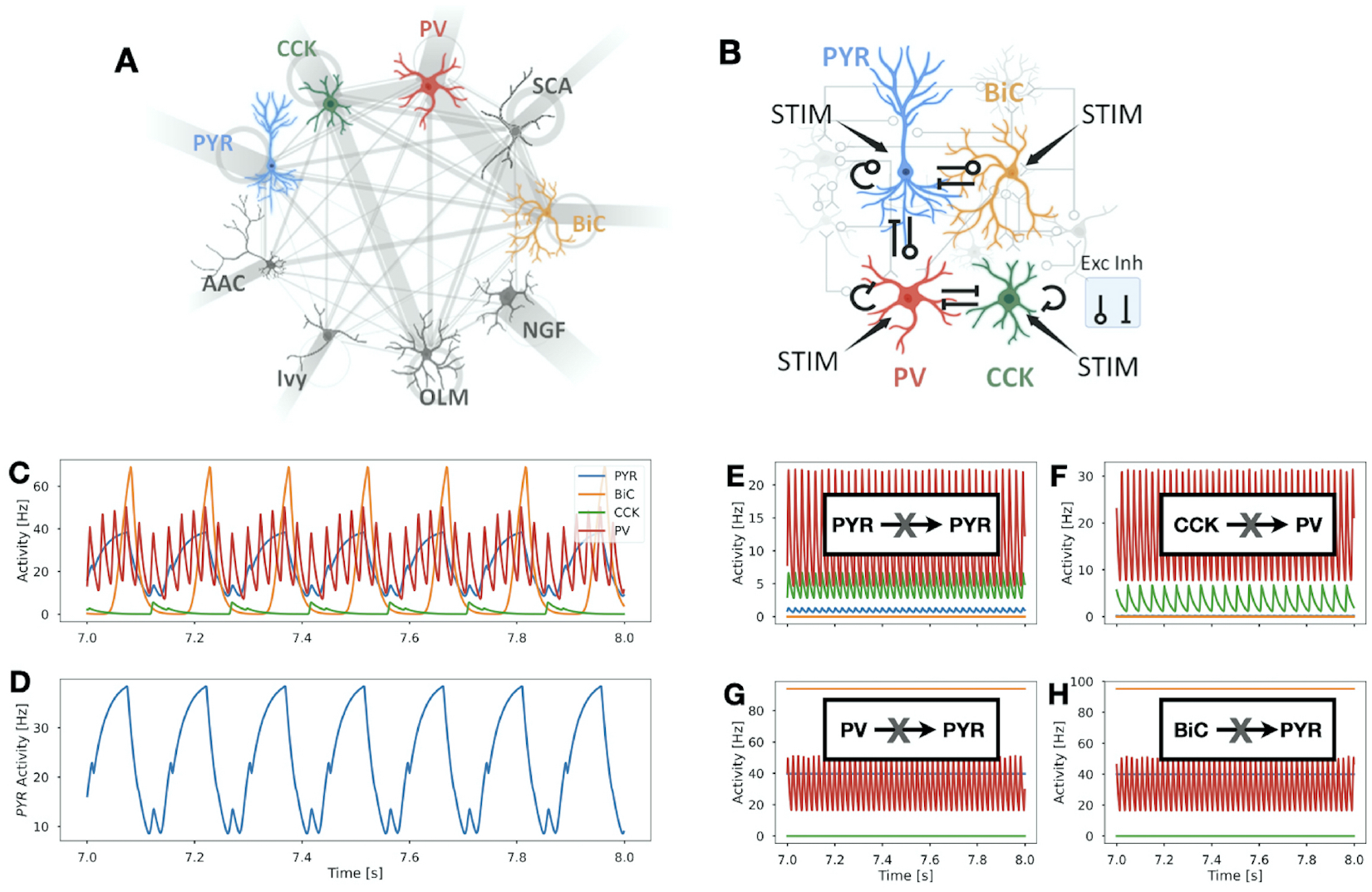
Schematics of microcircuit models of CA1 hippocampus and PRM output example. **A**. Stylized schematic showing the nine different cell types that are present in a detailed circuit model of CA1 hippocampus (Bezaire et al., 2016). These cell types are pyramidal cells, paravalbumin-expressing basket cells, axo-axonic cells, bistratified cells, cholecystokinin-expressing basket cells, Schaeffer Collateral-associated cells, oriens-lacunosum-moleculare cells, neurogliaform cells, and Ivy cells. They are referred to as PYR, PV, AAC, BiC, CCK, SCA, OLM, NGF, Ivy, respectively. Connections between the different cell types are represented by grey lines between them. Grey circles by a given cell type indicate that they are interconnected, and incoming grey lines to the different cell types represent external drives. The thickness of the lines and circles is representative of the relative strength of these various connections. **B**. Stylized schematic of the reduced model with four cell types (PYR, BiC, CCK, PV) and excitatory (Exc) and inhibitory (Inh) connections between them with possible inputs (STIM) to each cell type. The rationale used to consider this reduced model is given in the main text. **C**. Activity of the four cell types (PYR, BiC, CCK, PV) in the PRM at baseline (i.e., when no STIM is applied to any of the cell types) over a one second interval. Colors match those shown in the schematic of *B* of the four cell-type circuit. The synaptic weight parameter values used in this simulation are from *Set 4* in SUPPLEMENTARY TABLE 1. **D**. Activity of only PYR showing theta-gamma coupled rhythms. PYR output is analyzed as an LFP representation. **E-H**. Removal of connections between cell types as schematized in each panel. These connections were previously shown to be required for theta-gamma coupled rhythms. See text for further detail.

We built a PRM of the four cell microcircuit depicted in FIGURE 1 **B**, and explored it using a combination of techniques. Mathematical equations and parameter descriptions are provided in the Model and methods section. This PRM is novel relative to other mean field models in its incorporation of distinct cell types, particular interconnectivities and dynamics that speak of both theta and gamma oscillations and their co-existence. Its deliberate simplicity, compared to the other detailed models represents an important advantage as it enables a thorough exploration of parameter space with constrained values and a detailed mathematical characterization of different cell-type contributions. Since extracellular LFP recordings are typically done in the pyramidal cell layer with PYR constituting the vast majority of the cells in the network, we consider PYR activity as representing the LFP output. The PRM exhibits LFP oscillations within gamma and theta frequency bands that are coupled with one another. An example is shown in FIGURE 1**C**. We show the activities of all four cell types in FIGURE 1**C**, and only PYR in FIGURE 1**D** to more easily visualize the LFP representation of theta-gamma coupled rhythms.

**Table 1.** Parameter Values for PRM.

| Symbol | Definition | Value |
| --- | --- | --- |
| $\beta$ | Response function gain | 20 au |
| $\tau$ | delay | 5 ms |
| $\alpha_{PYR}$ | PYR rate constant | 40 Hz |
| $\alpha_{BiC}$ | BiC rate constant | 80 Hz |
| $\alpha_{CCK}$ | CCK rate constant | 40 Hz |
| $\alpha_{PV}$ | PV rate constant | 80 Hz |
| $ie_{PYR}$ | PYR intrinsic excitability | 0.03 au |
| $ie_{BiC}$ | BiC intrinsic excitability | -1.45 au |
| $ie_{CCK}$ | CCK intrinsic excitability | 0.8 au |
| $ie_{PV}$ | PV intrinsic excitability | 0.5 au |
| $r_{0,PYR}$ | PYR maximal firing rate | 40 Hz |
| $r_{0,BiC}$ | BiC maximal firing rate | 100 Hz |
| $r_{0,CCK}$ | CCK maximal firing rate | 60 Hz |
| $r_{0,PV}$ | PV maximal firing rate | 100 Hz |

The results are presented as follows. We first leverage a genetic algorithm to obtain constrained parameter sets. This reveals degeneracy in the expression of theta-gamma coupled rhythms and variability of the different connection types. That is, there are many different sets of synaptic weights that produce theta-gamma coupled rhythms. We then perform theoretical analyses on subsystems of the PRM. Use of the constrained parameter sets greatly reduces the dimensionality of the system, and allows us to obtain cellular insights into theta and gamma rhythm generation. We then carry out extensive numerical simulations and are able to expose contributions by the different cell types. In particular, we show that CCK exhibits multiple roles that include its ability to initiate the expression of theta-gamma coupled rhythms. In the last section, we bring all our results together and present essential cell-type specific motifs underlying the expression of theta-gamma coupled rhythms.

### Using a genetic algorithm exposes degeneracy in theta-gamma coupled rhythm expression

The co-existence of theta and gamma oscillations, as illustrated in FIGURE 1**D**, results from a combination of excitation-inhibition motifs, and delayed feedback. The frequency and power of these theta-gamma coupled rhythms are controlled by a combination of parameters. To determine which parameter combinations allow theta-gamma coupled rhythms in the PRM, we first applied constraints from the experimental literature for each of the four different cell types, while setting parameters to values that produce coupled oscillations within rationalized constraints (see Model and methods section). In doing this, we ensured that the overall dynamics of the network and relative differences in cell type properties were both taken into consideration.

One such important constraint is the manifest loss of theta activity following connection removal. Specifically, theta oscillations should no longer be present whenever connections between PYR and PYR, from CCK to PV, from PV to PYR, and from BiC to PYR, are removed (Chatzikalymniou et al., 2022). This constraint guided our parameter search: from an initial set of parameter values, we used a genetic algorithm search in the parameter space of the synaptic weights (*w*) and obtained 200 sets of synaptic weights satisfying the above criterion. For the example parameter set of FIGURE 1**C**, we show in **E-H** that the constraint of connection removals is satisfied.

The multiple sets of *w* obtained illustrate the degeneracy of the model system producing theta-gamma coupled rhythms. The distribution of these 200 sets of *w* is shown in FIGURE 2**A**. The variability differs for the nine different connection types, and this can be appreciated by observing their standard deviations (std) shown in the bar graph histogram of FIGURE 2**B**. Noting the large std for the synaptic weights involving CCK, we re-ran the genetic algorithm with an adjusted constraint regarding CCK. The outcome is shown in SUPPLEMENTARY FIGURE 2. The PYR→PYR connection had the largest synaptic weight for excitatory connections, and for inhibitory connections, the largest synaptic weights involved CCK: CCK→CCK and CCK→PV. These results start to suggest that CCK may have a more specific role relative to the other cell types.

**Figure 2.**
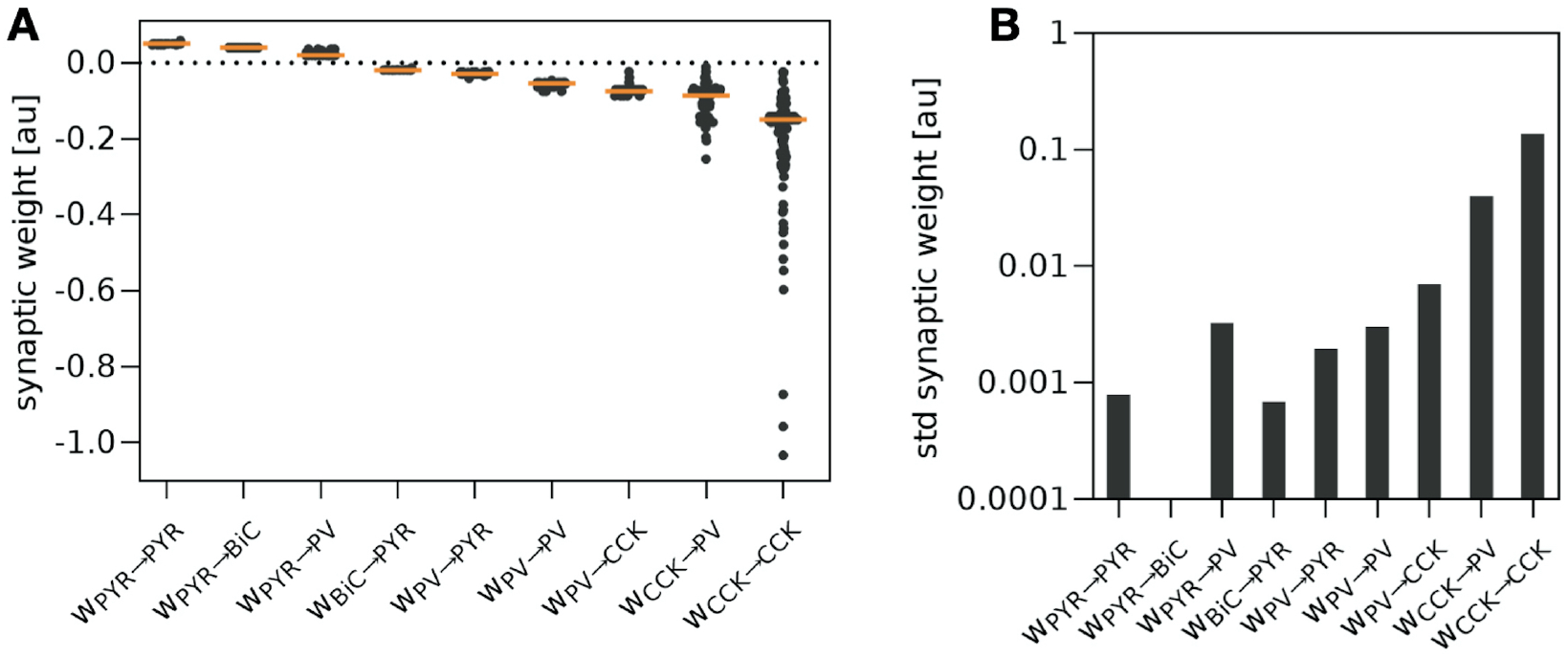
Distributions of constrained synaptic weights. **A**. Distribution of synaptic weights (*w*) for each of the nine different connections in the PRM, from the 200 constrained parameter sets of *w* found using a genetic algorithm. The orange line depicts the median. **B**. Bar graph showing the standard deviations (std) of *w* for each of the nine different connections.

From the 200 parameter sets shown in FIGURE 2**A**, we applied clustering techniques and selected ten parameter sets (see Model and methods section for details) for further analyses. These ten chosen parameter sets (*Sets 0-9*) represent a variety of synaptic weight parameter values that generate theta-gamma coupled rhythms in the PRM. Values of the synaptic weights for each of the nine different connections are given in SUPPLEMENTARY TABLE 1. With these ten constrained parameter sets in hand, we are able to greatly reduce the dimensionality of our theoretical analyses and to do extensive and systematic numerical explorations.

### Theoretical analyses predict circuit motifs underlying theta and gamma rhythms

To gain insights into the cell types and connectivity motifs contributing to the generation and interaction of theta and gamma rhythms, we leveraged linear stability analysis. We did this by systematically examining connectivity motifs responsible for generating oscillations in both the theta and gamma frequency bands. To achieve this, we dissected the PRM (see Eqs. (1a)-(1d) in Model and methods section) into three distinct and independent motifs — specifically PYR and BiC (PYR+BiC), PYR and PV (PYR+PV) and CCK and PV (CCK+PV) — each corresponding to a subsystem of dimension two. This approach enabled us to examine the behaviour of each motif, thereby enhancing our understanding of their contributions to theta and/or gamma activity. Additional details can be found in the Model and methods section. Activities of the cell types for the three subsystems as a function of time are shown in FIGURE 3**A, B, C**. The dynamics of PYR+BiC generates rhythms within the theta band, whereas both PYR+PV and CCK+PV motifs independently generate gamma frequency rhythms.

**Figure 3.**
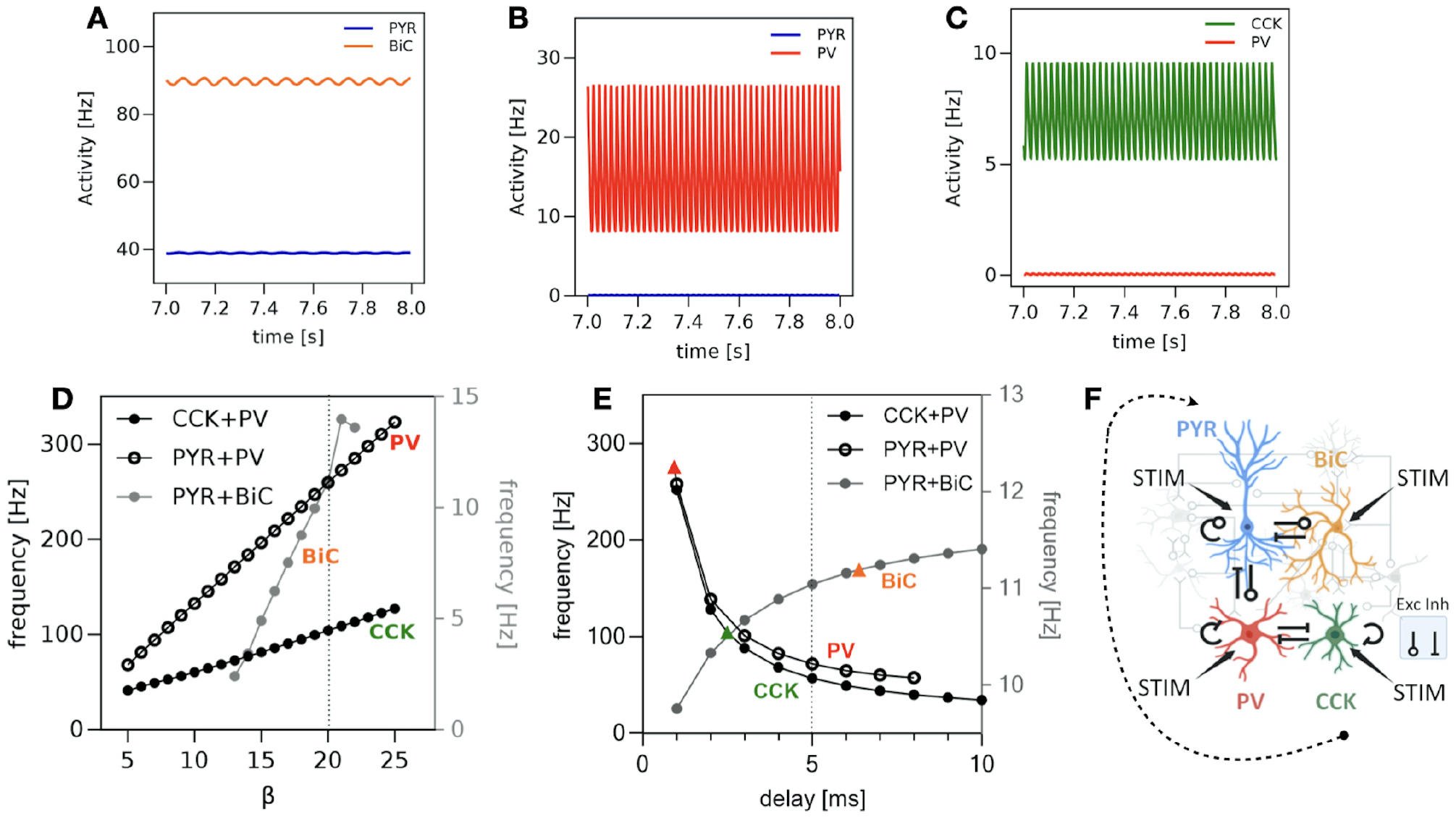
Theoretical analyses of PRM subsystems. **A**. Activity of the two cell types (PYR, BiC) in PYR+BiC subsystem over a one second interval showing theta rhythms. **B**. Activity of the two cell types (PYR, PV) in PYR+PV subsystem. PYR is silent while PV exhibits gamma frequencies. **C**. Activity of CCK and PV in the CCK+PV subsystem. CCK oscillates with gamma frequencies while PV demonstrates only a small amplitude and activity. **D**. Frequencies of BiC, CCK and PV are plotted for different values of *β*. **E**. Frequencies plotted for BiC, PV and CCK as a function of different delay values. The coloured triangles show the bifurcation points (that is, delays) for the three subsystems (PYR+BiC, PYR+PV and CCK+PV). In panels **D** and **E**, the frequencies are plotted for the cell type with the highest activity (between the two cell types of the subsystem). **F**. Schematic illustration of the inhibition mechanism of PYR initiated by CCK. *Set 0* parameter values were used in the calculations for these plots. In panels **A, B** and **C**, *β* = 20 and *τ* = 5ms.

While many parameters could have been considered, we focused our analysis on the roles of the response gain *β* as well as the synaptic and axonal time delay *τ*, given their well-known implication in promoting neural synchrony (Dhamala, Jirsa, & Ding, 2004; Erneux, 2009). We found that varying the value of *β* changes the frequencies of the different cell types. Specifically, frequency increases with increasing *β* (FIGURE 3**D**). This relation comes directly from the stability analysis (see Eqs. (16a)-(16b) in Model and methods section). The time delay was found to influence the oscillatory behavior of each subsystem. To study its role in the generation of theta and gamma activity, we fix *β* = 20 and further analyze the frequencies of each of the three subsystems. In FIGURE 3**E**, the frequencies of the three subsystems are depicted as a function of delays. For each subsystem, we plotted the activity of only one cell type because either the second cell type remains silent or exhibits a very small amplitude (see FIGURE 3**A-C**). Notably, the frequency of PYR+BiC consistently remains in the theta band across all selected delay values. In contrast, CCK+PV and PYR+PV subsystems demonstrate higher frequencies than gamma for small delays. As delays increase, the associated frequency values decrease, ultimately settling within the gamma band. The linear stability analysis further reveals that these oscillations emerge through Hopf bifurcations. To maintain simplicity in our model, for the remainder of the study, we fix the delay at a value at which our analysis predicts the co-existence of both theta and gamma frequencies. We choose five milliseconds as this is reasonable considering physiological constraints (see Model and methods).

The activation of CCK in the CCK+PV subsystem (FIGURE 3**C**) and PV in the PYR+PV subsystem (FIGURE 3**B**) suggest a potential relationship between the initiation of gamma frequencies and the inhibition of PYR through a mechanism where CCK is implicated in the initiation of gamma frequencies and PV subsequently when it inhibits PYR cells. This is schematized in FIGURE 3**F**. However, the general result holds for all of the ten parameter sets (see parameter values in SUPPLEMENTARY TABLE 1).

### Numerical explorations support theoretical predictions for circuit motifs underlying theta and gamma rhythm generation

To assess the susceptibility of each cell type to the expression of theta-gamma coupled rhythms, we systematically explored our four-cell circuit by applying a stimulus (i.e., STIM, see FIGURE 1**B**) to each of the four different cell types, using the ten chosen parameter sets (SUPPLEMENTARY TABLE 1). We performed simulations for a wide range of STIM values and measured stimulation ranges over which theta-gamma coupled rhythms persist to assess the robustness of the coupling and the roles played by the different cell types in influencing their existence.

The stability analysis predicted that PYR+BiC circuits can function as theta frequency rhythm generators, but not any of the other two cell subsystems. For gamma rhythms, either PYR+PV or CCK+PV (or both) lead to gamma frequency rhythm expression. To consider this in the full four-cell circuit, we examined STIM values which preserved the theta-gamma regime and determined how sensitive the spectral properties of the co-existing oscillations were to selective stimulation. We considered theta-gamma coupled rhythms to be preserved (i.e., sufficiently present) if both theta and gamma rhythms had powers that were at least 25% of a reference when STIM was zero (baseline). We quantified theta and gamma powers from the theta-gamma coupled rhythms, and calculated the slope of a linear fit to theta or gamma power plots with STIM. This is illustrated in SUPPLEMENTARY FIGURE 3. In FIGURE 4, we plot the values of these fitted slopes for each cell type for theta power (**A**), and gamma power (**B**).

**Figure 4.**
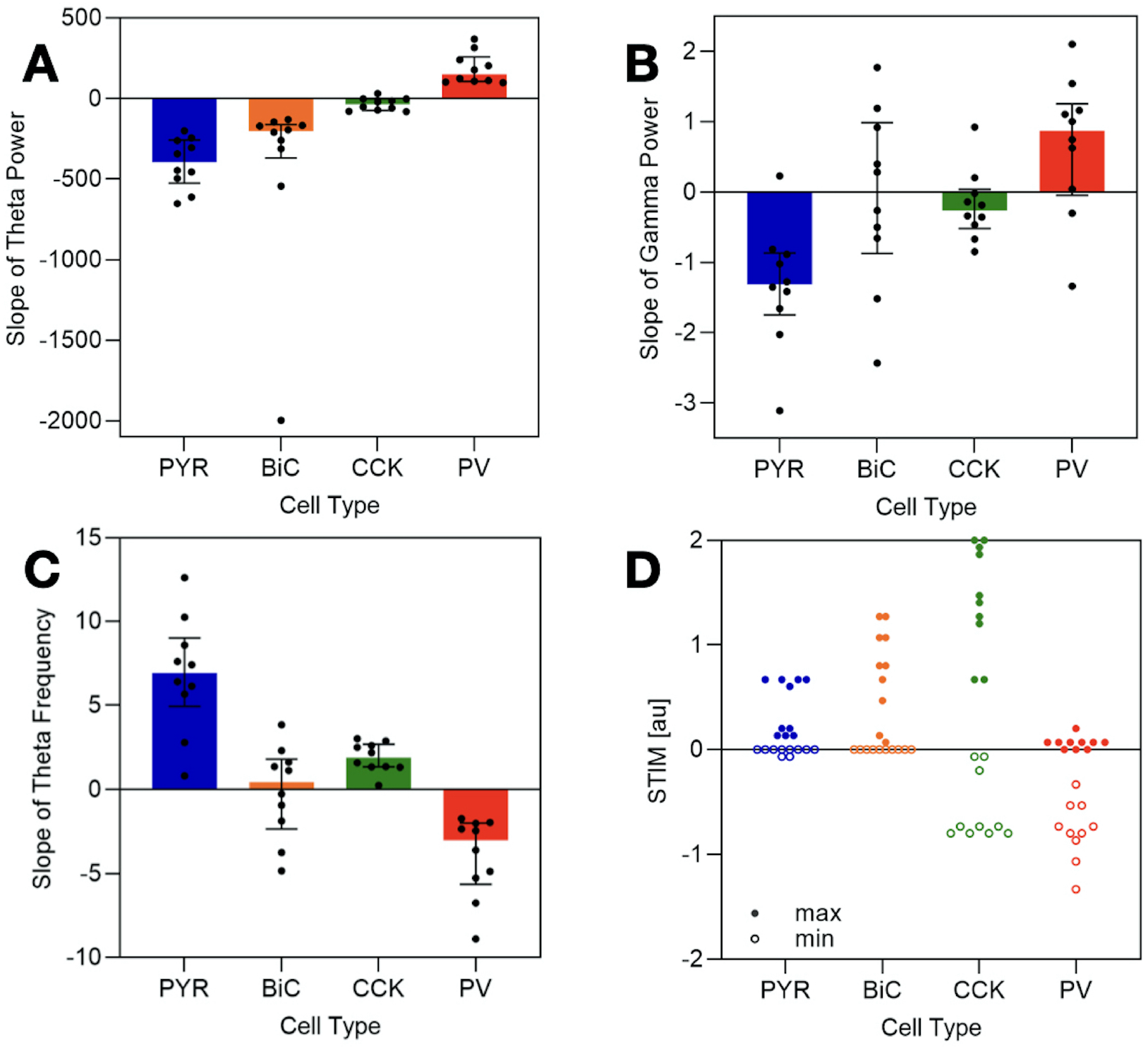
Cell type sensitivities. STIM values are applied to each cell type for each of the ten parameter sets (*SUPPLEMENTARY TABLE 1*). Theta and gamma powers and theta frequency are extracted when theta-gamma coupled rhythms are present. Plots show the distribution of the slope of the theta power (**A**), gamma power (**B**) and theta frequency (**C**) with increasing STIM to each cell type. The slope is derived from a linear best-fit (see *SUPPLEMENTARY FIGURE 3*) through the points where the system exhibits theta-gamma coupled rhythms (see Model and methods section for further details). For all of the plots in *A-C*, the bars extend from the first quartile (Q1) to the third quartile (Q3) of the data, shown with the median. The colors are representative of the particular cell type (see FIGURE 1B). **D**. Minimum (min) and maximum (max) STIM values to the different cell types that allowed theta-gamma coupled rhythms to be expressed. Statistical significances are provided in *SUPPLEMENTARY TABLES 4 and 5*.

We note that theta power increases (i.e., positive values in FIGURE 4**A**) existed only with increasing STIM to PV. PYR and BiC both showed theta power decreases and there was minimal effect with CCK. Considering their medians, maximal absolute changes occurred for PYR and BiC, although BiC showed a large variability. This is interesting since our theoretical analyses above predicted that only PYR+BiC circuits, and not the other two-cell circuit subsystems, produce theta frequency rhythms (see FIGURE 3**E**). Given that our numerical explorations showed that the theta power changed maximally for PYR and BiC when they were stimulated indicates that the magnitude of theta power change with stimulation in our numerical explorations of the four-cell system is consistent with the expression of theta rhythms in our theoretical analyses of the subsystems. For gamma power, the maximal absolute changes occurred for PYR and PV, as shown in FIGURE 4**B**. Given the consistency between the numerics and the stability analysis outcome for theta rhythms, in a similar vein, our numerical exploration observations suggest that the PYR+PV subsystem could be more key for gamma rhythm generation relative to the CCK+PV subsystem. However, as the gamma powers are much smaller than the theta powers (see SUPPLEMENTARY FIGURES 3 and 4), more investigation is needed to extract further particular contributions of cell types for the fast rhythms.

Given the functional importance of theta frequencies, we examined how they changed, as part of the theta-gamma rhythms, when the different cell types were stimulated. In FIGURE 4**C**, we show a quantification of how much theta frequency changed and note that the largest absolute changes occurred with PYR and PV stimulation. BiC showed both increases and decreases in frequency with stimulation.

### CCK plays multiple roles in theta-gamma coupled rhythm expression

We measured the range of STIM values associated with sufficient theta-gamma coupling. In FIGURE 4**D**, we present the minimum and maximum STIM values for each cell type. It is evident that CCK exhibits the widest range, indicating that theta-gamma coupled rhythms can be present for a wider range of external inputs compared to the other cell types in the network. PYR has the smallest range (see SUPPLEMENTARY TABLE 4 for statistical significance).

Given the larger synaptic weights and variability involving CCK, as exposed by use of a genetic algorithm, we thought that CCK may have more specific roles compared to other cell types. We thus carried out additional investigations that revealed multiple CCK contributions. We removed CCK by silencing them using a strong inhibitory stimulus, either at the beginning (*middle plots* of FIGURE 5) of the simulation or during (*top plots* of FIGURE 5) ongoing theta-gamma coupled rhythms in the ten sets. Removal at the start of the simulation *(middle plots)* consistently prevented theta-gamma coupling — note that all cell types begin in a quiescent state across all simulations (see Model and methods section). Introducing a CCK ‘burst’ to the circuit, previously devoid of theta-gamma coupled rhythms due to the absence of CCK, resulted in the expression of theta-gamma coupling (*middle plots*). Upon examining the timing of CCK and PV following a CCK ‘burst,’ we found that CCK firing precedes PV firing for all ten sets (not shown). These results support the notion that CCK is central in initiating the expression of theta-gamma coupled rhythms, as suggested from our theoretical analyses (see FIGURE 3**F**). However, once coupled rhythms are expressed, CCK activity is not necessarily required. In *Sets 3, 5* (shown in FIGURE 5**B**), and *8*, where CCK activity was needed for the continuation of theta-gamma coupled rhythms (*top plot*), we observe larger connections between PYR and PV (see SUPPLEMENTARY TABLE 1) suggesting that a sufficiently active PYR can sustain theta-gamma coupled rhythms without CCK activity in the circuit.

**Figure 5.**
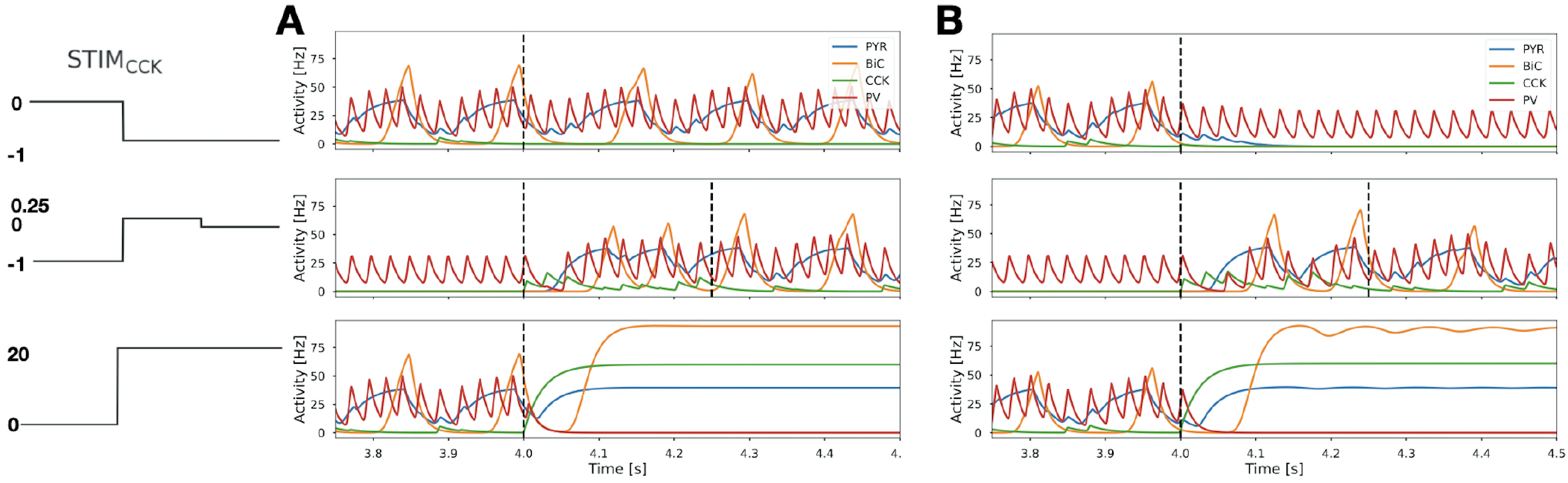
Effects of CCK perturbation on theta-gamma coupled rhythms. (*Top plots*) CCK is removed from the circuit at 4.0 seconds by changing STIM_*CCK*_ to -1 (vertical dashed line) leading to continuation of theta-gamma coupled rhythms for *Set 4* (**A**), or to loss of coupled rhythms for *Set 5* (**B**). (*Middle plots*) CCK is removed from the circuit at the start of the simulation, and at 4.0 seconds, CCK is ‘added back’ to the circuit by changing STIM_*CCK*_ to 0.25 for a brief interval (as delineated by the first vertical dashed lines), and then returned to baseline (i.e., STIM_*CCK*_ to 0) at the second vertical dashed line. Theta-gamma coupled rhythms are expressed when CCK is added back to the circuit. (*Bottom plots*) CCK activity is strongly enhanced in the circuit at 4.0 seconds by changing STIM_*CCK*_ to 20 (vertical dashed line) leading to loss of theta-gamma coupled rhythms for *Set 4* (**A**), or *Set 5* (**B**). Representation of the changes in STIM_*CCK*_ are shown on the left hand side for each row of plots.

We further examined whether there would be any change in the regularity of the ongoing theta rhythm with or without CCK during theta-gamma coupled rhythms. Analyzing intervals between PYR peaks (i.e., the cycle period of the LFP representation), we observed a notable increase in the standard deviation of these intervals when CCK was absent from the circuit producing theta-gamma coupled rhythms in the majority of sets (see SUPPLEMENTARY TABLE 3 and SUPPLEMENTARY FIGURE 5). This suggests that CCK could play a role in regularizing the spectral features of theta activity.

Lastly, strong activation of CCK silences PV, preventing coupled rhythms as gamma rhythms are no longer present in PV and they cannot be propagated to PYR from CCK (see *bottom plots* of FIGURE 5). Thus, the level of CCK activity, along with the activity of the other cells in the circuit, can serve as a determinant for the expression of theta-gamma coupled rhythms, suggesting a potential ‘switching’ role for CCK.

### Bringing it all together

Due to the combined insights from our theoretical analyses and numerical explorations of the PRM, we can assign particular roles to the different cell types.

PYR and BiC play the role of theta rhythm generators. This is schematized in FIGURE 6**A**. This is based on the theoretical analyses of the subsystems in which only the PYR+BiC subsystem led to rhythms in the theta frequency band as fully supported by the numerical simulations of the full four-cell system that showed that PYR and BiC more strongly affected theta power relative to the other cell types. CCK contributes to the gamma rhythm portion of theta-gamma coupled rhythms, serving as ‘initiators’ of the gamma rhythm (see FIGURE 5, with PV contributing to the gamma by ‘applying’ it to PYR. This is schematized in FIGURE 6**B**. Gamma rhythms can emerge from PYR+PV or CCK+PV subsystems. PV acts as an ‘enhancer’ of both theta and gamma. That is, it is the only cell type that leads to an increase in *both* theta and gamma power with increasing stimulation (see FIGURE 4**A**,**B**).

**Figure 6.**
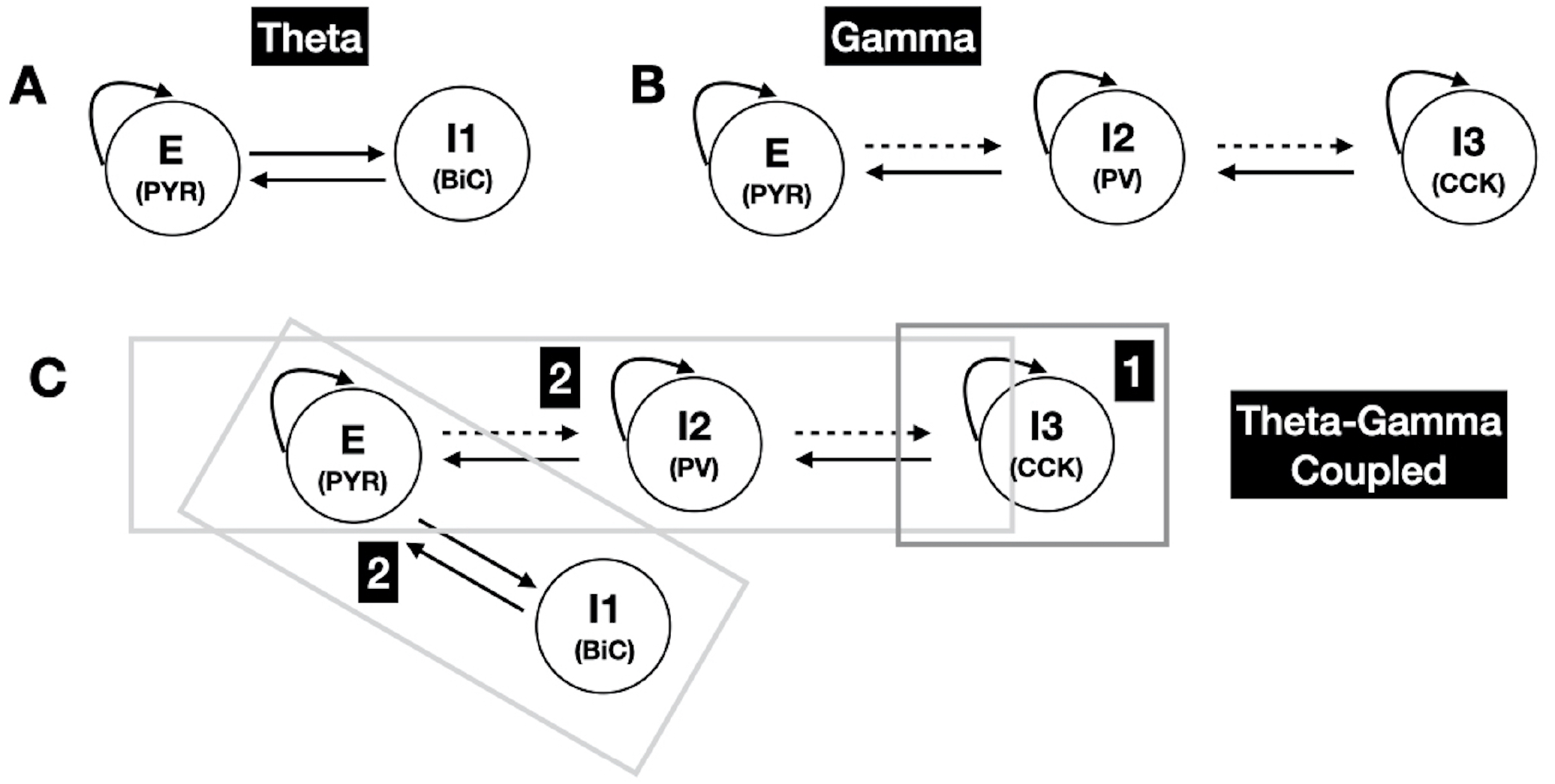
Predicted motifs for coupled rhythms. As derived from theoretical and numerical analyses of the four-cell circuit shown in FIGURE 1*B* with ***E***=PYR, ***I1***=BiC, ***I2***=PV, ***I3***=CCK. **A** shows the motif for theta rhythm generation, **B** for gamma rhythm generation, and **C** shows the four cell motif for theta-gamma coupled rhythm generation. It occurs in two phases as shown in *C*. In ***phase 1, I3 (CCK)*** initiates the coupled rhythm by being able to disinhibit ***E (PYR)***, which then allows theta rhythms to be generated (via ***E (PYR)*** and ***I1 (BiC)***) together with gamma rhythms (via ***E (PYR), I1 (BiC)*** and ***I2 (PV)***) in ***phase 2***. The dashed connection lines means that they are not critical for the system to be able to express theta-gamma coupled rhythms — see SUPPLEMENTARY TABLE 2. These motifs imply that at least three different inhibitory cell types (I1, I2, I3) need to be part of the circuitry for coupled rhythms to be expressed from quiescent states.

Given the described roles for the different cell types, we expect that when the PV→PYR connection is removed, there would be no gamma (as it is coming from PV and CCK via PV), and thus no theta-gamma coupled rhythms. However, with removal of the PYR→PV connection type, we expect that theta-gamma coupled rhythms might still be possible as theta could still exist due to the PYR+BiC subsystem, and gamma could still ‘circulate’, as schematized in FIGURE 3**F**. In SUPPLEMENTARY TABLE 2, we see that theta-gamma coupled rhythms can be present for some of the parameter sets. Theta-gamma coupled rhythms can remain when the PV→CCK connection is removed (see SUPPLEMENTARY TABLE 2), which makes sense since CCK can initiate the coupled rhythms. If PYR, PV and BiC cells are quiescent, or not active enough, CCK is needed to initiate theta-gamma coupled rhythms (see *middle plots* of FIGURE 5). But it is possible that CCK is not needed to maintain theta-gamma coupled rhythms if the different cells’ histories are such that they are active enough (see *top plot* of FIGURE 5**A**). However, CCK clearly contributes to the regularity of the theta rhythm (see SUPPLEMENTARY TABLE 3).

We thus present a two-phase process for the generation of theta-gamma coupled rhythms in the hippocampus, as schematized in FIGURE 6**C**. In the first phase, CCK generates gamma rhythms to initiate the theta-gamma coupled rhythm process. It can do this prior to PV contributing to gamma rhythms in light of its intrinsic properties (higher input resistance) and its surrounding milieu (see SUPPLEMENTARY FIGURE 1). SUPPLEMENTARY FIGURE 6 shows examples from two of the parameter sets illustrating CCK peaking before PV. In the second phase, theta is generated (by PYR and BiC) and gamma is enhanced (by PYR and PV), leading to theta-gamma coupled rhythms. In essence, the disinhibitory effect of CCK to PYR plays a controlling role in the two-phase process. Overall, we suggest that the mechanisms portrayed in FIGURE 6**C** underlie theta-gamma coupled rhythms, and could serve as a motif ‘template’ for the existence of coupled rhythms of high and low frequencies in the brain.

## DISCUSSION

### Summary and predictions

Combining theoretical and numerical insights of a four-cell circuit model of CA1 hippocampus, we have presented a mechanism for how theta-gamma coupled rhythms arise, showcasing the contribution of different inhibitory cell types. Using a mean field model with distinct cell types, our analyses allowed us to present circuit motifs underlying the co-expression of high and low frequency rhythms. Gamma rhythms in theta-gamma co-expression involve a three-cell circuit motif (FIGURE 6). For only gamma, the theoretical work pointed to either PYR and PV or CCK and PV circuits as the gamma generators. Further, removal of PV→PV or PV→PYR connections abolished gamma rhythms, indicating support for either PING or ING type mechanisms (Whittington et al., 2000).

From our systematic numerical explorations together with theoretical analyses, we predicted that PYR and BiC underlie theta rhythm generation as they showed the largest modulation in theta power when stimulated (see FIGURE 4**A**). We had previously postulated that an ‘inhibition-based tuning’ was responsible for the generation of theta rhythms (Skinner, Rich, Lunyov, Lefebvre, & Chatzikalymniou, 2021), and our work here predicts that BiC are the inhibitory cell type that can initiate this generation. PYR and PV showed the most modulation when theta frequency was considered (see FIGURE 4**C**). This is consistent with our previous work using detailed circuit models where we had found that inputs to PYR were able to set the population theta rhythm frequency, increasing with larger net excitatory drive (Chatzikalymniou et al., 2021). Here, with our PRM, we found that theta frequency is most strongly affected by PYR stimulation, increasing when PYR were stimulated, that is, an increased net excitatory drive

It is known that CCK is a cell type that is highly modulated (Freund & Katona, 2007). In particular, it is known that CCK expresses highly asynchronous postsynaptic potentials (Daw, Tricoire, Erdelyi, Szabo, & McBain, 2009) making it notable that we found that the CCK→PV connection type is more variable relative to other connection types for theta-gamma coupled rhythms (see FIGURE 2**B**). From our theoretical work using two-cell subsystems, we had suggested that CCK initiated gamma rhythms, and from our numerical explorations, we found that CCK is needed to initiate theta-gamma coupled rhythms from quiescent states. Recent experimental studies have shown alternating sources of perisomatic inhibition onto pyramidal cells during behaviour via CCK-expressing and PV-expressing basket cells involving inhibitory mechanisms between these cell types (Dudok et al., 2021). Theta-gamma coupled rhythms are expressed during movement exploration and REM sleep (Buzsaki, 2011), and thus, it would be interesting to examine whether these coupled rhythms are differentially affected by perturbing either PV-expressing or CCK-expressing basket cells, and whether CCK-expressing basket cells play a more robust role relative to other inhibitory cell types in the expression of theta-gamma coupled rhythms. Our results here predict that the removal of CCK-expressing basket cells should drastically reduce the presence of theta-gamma coupled rhythms, although perhaps not eradicate it, and the regularity of the theta rhythms should be reduced. Further, we predict that modulation of CCK-expressing basket cells could lead to theta-gamma coupled rhythms being turned on or off, thus acting to control the ‘mood’ (as borrowed from Freund (2003) terminology) of the system.

### Related studies

A recent data-driven modeling study probed theta-gamma phase amplitude coupling (PAC) in hippocampal circuits focusing on three cell types - pyramidal cells, PV-expressing basket cells and OLM cells (Ponzi, Dura-Bernal, & Migliore, 2023). Multi-compartment detailed cellular models were used along with short-term synaptic plasticity, allowing the authors to highlight these cell types and particular aspects. Their results indicated a dependence of theta oscillation expression on gamma for the ubiquity of PAC output via these cell types and their interconnections. Specifically, they showed that the theta rhythm frequency and strength relied on a PING mechanism. Our PRM is supportive of PING-like mechanisms and is thus consistent with Ponzi et al. (2023). However, due to the limited nature of our LFP representation, we did not apply PAC measures to them in the PRM.

Very detailed network models with explicitly identified and characterized cell types have been developed and can express theta and gamma rhythms (Bezaire et al., 2016; Ecker et al., 2020; Romani et al., 2024). Although it is not possible to parse mechanisms using these models, they can be used to explore biological intricacies such as ion channel complements in different cell types and receptor densities. Theta-gamma oscillations during neurostimulation has recently been explored (Vardalakis, Aussel, Rougier, & Wagner, 2024), but the theta rhythms were imposed on the hippocampal system. For experimental linkage design considerations, it may be interesting to explore neurostimulation protocols with the PRM generating theta-gamma coupled rhythms.

### Limitations and future work

Several biological aspects are not directly represented in the PRM. This includes aspects such as spatial inhibitory distributions and inputs onto pyramidal cells, short-term plasticity as well as the inclusion of additional inhibitory cell types. The results from our study do not of course imply that these other aspects are not important. Rather, our results are limited by the cell types and interconnections that were included (FIGURE 1**B**). It may be possible to use our existing PRM as a starting base to consider additional cell type contributions. For example, the contribution of OLM cells could be initially considered in an indirect way through its effect on bistratified cells by considering a decreasing STIM (inhibitory input) onto them - the PRM would indicate an enhanced theta power (see FIGURE 4**A**). Specific known motifs that include VIP-expressing cell types (Guet-McCreight, Skinner, & Topolnik, 2020) can also be considered indirectly. Interneuron-specific 3 cells (VIP-expressing subtype) are known to differentially inhibit OLM cells, bistratified and basket cells (Tyan et al., 2014), and so an initial consideration of inputs received by these different cell types and their effect on theta-gamma coupled rhythms could be probed. Also, as theta frequency and behavioural correlates are known to vary across the dorsoventral axis (Strange, Witter, Lein, & Moser, 2014), we can consider leveraging our models to understand these differences from particular cell types and interconnection variations. In general, one could envision expanding the PRM to directly include additional cell type populations and connectivities and interpretations to develop further hypotheses and obtain predictions for experimental examination.

The relative strength of connections between inhibitory populations seems to change across the septo-temporal axis of the CA1 hippocampus and across layers (Navas-Olive et al., 2020; Soltesz & Losonczy, 2018). Navas-Olive et al. (2020) have shown that PV-expressing basket cells preferentially innervate pyramidal cells at the deep sublayers while CCK-expressing basket cells are more likely to target the superficial pyramidal cells. Also, there is different innervation of pyramidal cells by bistratified and OLM cells for deep versus superficial pyramidal cells. We can start to consider how these differential connectivities affect theta-gamma coupled rhythm control by focusing on such perturbations in the PRM. Interestingly, we note that when connections from PV-expressing basket cells to pyramidal cells are larger (smaller), there can be a smaller (larger) theta frequency (see SUPPLEMENTARY TABLE 1, last column), suggesting theta frequency differences between deep and superficial layers due to particular connection weight differences.

For the PRM used here, despite our experimental linkage of the different cell type’s characterization, further examination of the different parameters is warranted. For example, different and differential *β*’s and delays could be considered to better capture the different cell types, and the formal inclusion of synaptic time constants could be considered, as well as differential milieu and intrinsic excitabilities. Additional theoretical analyses may be possible, but here we were able to exploit stability and bifurcation analyses to extract our insights.

## MODEL AND METHODS

### The population rate model (PRM)

The purpose of the PRM is the identification and analysis of key mechanisms involved in the generation and control of theta-gamma coupled rhythms, that is, coupling interactions between theta and gamma rhythms, in hippocampal microcircuits, following the formalism of population-scale mean-field models. The PRM describes the time evolution of PYR, BiC, CCK, and PV activities or mean firing rates (*r*_*PY R*_, *r*_*BiC*_, *r*_*CCK*_ and *r*_*PV*_) as a function of their mutual connectivity motif (FIGURE 1**B**), synaptic weights (*w*_*n*→*m*_, for *n, m* = PYR, BiC, CCK, PV), membrane rate constants (*α*_*PY R*_, *α*_*BiC*_, *α*_*CCK*_ and *α*_*PV*_), synaptic and axonal delays lumped in a single term (*τ*), cell-specific intrinsic excitability (*ie*), and how these collectively contribute to the generation of rhythmic firing rate expression. The resulting model builds on the well-known (H. R. Wilson & Cowan, 1972) formalism, resulting in the following set of nonlinear, delayed differential equations,

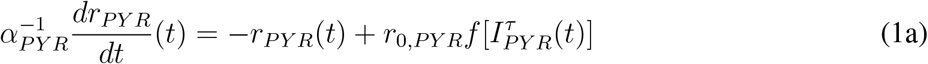

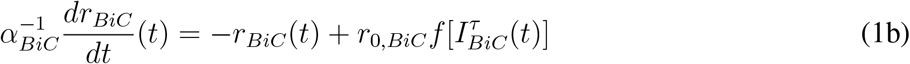

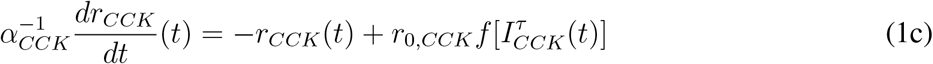

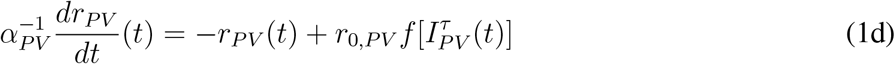

where *r*_0,*m*_ (here and below, *m* = PYR, BiC, CCK or PV) represents the maximal firing rate of the neurons. Fluctuations in mean firing rates result from the nonlinear integration of presynaptic inputs *I*_*m*_. Presynaptic inputs *I*_*m*_ scale firing rates through the firing rate response function

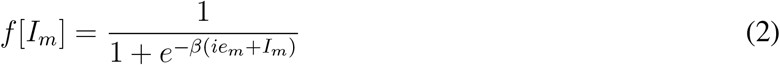

whose gain *β* reflects the steepness of the response, and intrinsic excitability (*ie*_*m*_) allows cell-specific threshold, rheobase, and input resistances to be taken into consideration. The presynaptic inputs for the four cell types considered are given by

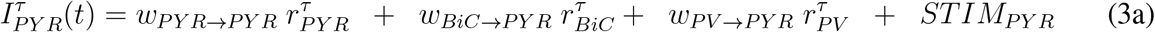

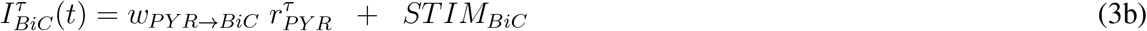

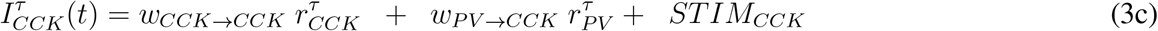

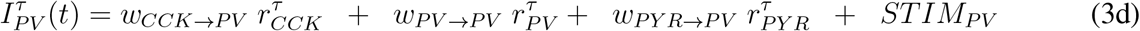

where 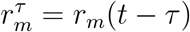. A time delay of *τ* = 5 ms is included to represent physiological axon conduction delays and synaptic activation time values. The *α*_*m*_ parameter sets the time scale at which populations respond to stimuli. Cell-specific sensitivities and control are explored by including an additional input (*STIM*_*m*_).

### Choosing the initial PRM parameter set

Based on the experimental literature, we set parameter values for membrane rate constants (*α*’s) and maximal firing rates (*r*_*o*_’s) of the different cell types. For the numerical explorations, *β* was fixed and set to the same value for all cell types, and *τ* was set to five milliseconds as a reasonable value considering physiological axon conduction delays and synaptic activation time values (Traub, Miles, & Buzsáki, 1992). We determined *ie*’s for the different cell types and a set of synaptic weights (see TABLE 1 and see *Set 0* in SUPPLEMENTARY TABLE 1) that allowed the system to produce theta-gamma coupled rhythms. The experimental literature resources were derived from online databases (Wheeler et al., 2015) and Table 6 and Appendix 1 of (Bezaire et al., 2016). The resulting ‘intrinsic’ activities (i.e., when all synaptic weights are zero) are shown in SUPPLEMENTARY FIGURE 1, where the different maximal firings and differential STIM responses (considering rheobase and input resistance values) are consistent cellular correlates in the literature. We note that zero synaptic weights here do not prevent an interpretation that the different cell types receive inputs from other sources, just not from the other cell types included here. That is, the interpretation of a given cell’s activity when disconnected from the other cell types in the circuit (SUPPLEMENTARY FIGURE 1) is that its activity represents its intrinsic behaviour in an existing milieu that includes inputs from any other cells not directly represented in the four-cell circuit system.

### Simulations

Simulations of the PRM were performed using an Euler method to numerically integrate the differential equations with a time-step of 0.001 seconds. Each simulation was performed for 8.0 seconds. The initial conditions were set to zero for all cell types. The first second of the simulations was not used in the spectral analysis to avoid the transients at the start of the simulations from affecting the analysis. The output of these simulations contains the activity of the four cell types in the model. These represent the population firing rate for the corresponding cell type.

### Frequency bands and power spectral density (PSD)

We define the low-frequency *theta* band to be between 3-12 Hz and the high-frequency *gamma* band to be between 20-100 Hz. We used a Welch periodogram of the signal and separated it into the two frequency bands. The dominant frequency and its corresponding power were calculated as the frequency where the highest peak appears in each frequency band and the power spectral density (PSD) at that frequency.

We found that filtering the signal into the low and high frequency bands caused anomalies due to the harmonics of the theta signal having a higher power than the gamma signal for PYR. This is due to the large difference in power between the theta and gamma bands in PYR activity. To circumvent the issue where the harmonics of PYR theta activity were stronger than PYR gamma activity, we used the periodogram of PV activity to find where the dominant frequency in the gamma band was present. This was done because PV activity has a stronger gamma component and as such, the harmonics of its activity in the theta band do not produce spurious results. Using the dominant frequency in the gamma band that we derived from the periodogram of PV activity, we find the corresponding PSD for PYR at that frequency to calculate the PSD in the gamma band.

We interpret PYR as an analog of the LFP because of the much larger prevalence of PYR compared to all the other cells in the network in actual brain tissue and since LFP recordings are mostly done in the pyramidal cell layer where PYR cell bodies are located. The average theta and gamma powers from the ten selected parameter sets (see SUPPLEMENTARY TABLE 1) were used as a reference to determine whether simulation outputs had theta and gamma rhythms of a considerable enough power. Oscillations were considered ‘sufficient’ if both the theta and gamma power were at least 25% of this reference. Considering both theta and gamma powers in this way ensured that co-expression was reasonably present, as would be reflected in a ratio of the two powers. Performing more sophisticated analyses of cross-frequency coupling (CFC) was not deemed appropriate for these model LFP signals as the type of CFC could take many forms such as phase-amplitude coupling (PAC), phase-phase or amplitude-amplitude coupling, and details of the signal shape, for example, could produce many confounds (Scheffer-Teixeira & Tort, 2022).

### Hypotheses testing and genetic algorithm

Using the initial set of synaptic weights (*Set 0*) to start, a genetic algorithm was used to explore the parameter space of the nine synaptic weights and find additional sets of synaptic weights that would produce theta-gamma coupled rhythms satisfying our hypotheses and experimental constraints.

A given set of synaptic weights must satisfy the following constraints:

- Primary: At baseline (i.e., no connections removed, and no STIM applied), the system should exhibit theta and gamma oscillations such that:
  - the power of PYR at the most prominent frequency in the theta and gamma bands are at least 60% of that in *Set 0*.
  - the ratio of the maximal firing rate (i.e., activity) for PV and BiC is at least 0.67
  - the maximal firing rate of CCK is greater than 4
- Secondary: When a certain connection type is removed, the theta oscillations break down (Chatzikalymniou et al., 2022).
  - These connection types are: PYR→PYR, CCK→PV, PV→PYR, and BiC→PYR.
  - This is quantified by the power of the most prominent frequency in the theta band for PYR being less than 10% of that in *Set 0* at baseline.

Using our genetic algorithm, we ‘mutated’ the synaptic weights of a constrained parameter set and checked if the resultant new set satisfied our hypotheses. If a new set does satisfy all the hypotheses, it is added to the list of the constrained sets for the next iteration of mutations. This was continued until a target number of constrained sets was found — that we set to be 200 — or a maximum number of iterations were completed

The resulting parameter sets were then clustered in the *w* parameter space using a *k-means* clustering algorithm (Pedregosa et al., 2011). We found 13 distinct clusters from the 200 parameter sets. To reduce this, we decided to only analyze the clusters where at least one set of synaptic weights produced oscillations where the ratio of the activities of PV and BiC was at least 0.70 — that is, increased from 0.67 used in the primary constraints. This ratio is based on relative firing rates presented in Table 6 of Bezaire et al. (2016), and the firing rate for CCK was chosen to ensure its participation in the overall system (as noted from multiple simulations). Using this additional threshold, we discarded three of the clusters that we had found. The ratios and CCK activities for these sets are given in SUPPLEMENTARY TABLE 1.

### Varying STIM

To explore how each cell type would respond when perturbed, stimulation (STIM) applied to each cell type was varied as a form of sensitivity analysis. Specifically, the STIM to each cell type was varied between -2 and 2, keeping the STIM to all other cell types fixed at zero. A positive value of STIM would mean that a cell type was being excited more than at baseline (where no STIM is applied) while a negative value of STIM would mean that the cell type is inhibited more than at baseline.

We used 121 equally spaced points in the range of values we explored. At each STIM value, the PRM was simulated and the theta frequency, theta power, and gamma power of PYR activity (i.e., LFP output representation) was calculated. A line of best fit was calculated for each of theta frequency, theta power and gamma power to quantify their change with respect to STIM to a particular cell type (see SUPPLEMENTARY FIGURES 3 and 4). The STIM range was considered as the difference between the largest and smallest STIM values that produced sufficient oscillations. Only points that showed sufficient oscillations (see above) were used for these calculations.

### Testing for regularity

The system was simulated for a total of ten seconds, with the same time-step as previously used. For the first five seconds, STIM to all cell types was set to zero. At t = 5 seconds, STIM_*CCK*_ = -1 was applied to silence CCK. The recording was split into two sections of four seconds each: the section from t = 1-5 seconds when CCK is active and the section from t = 6-10 seconds when CCK is silent. The first one second interval after the start of the simulation and after STIM_*CCK*_ = -1 was applied is ignored to not count any transient activity in the analysis. For each section, the peaks in the PYR and BiC activity were recorded. The mean, median, and standard deviation of the amplitudes of the peaks and the inter-spike interval (ISI) were recorded. This procedure was performed for all parameter sets where applicable.

### Linear stability analysis

For the analysis we switch to the following notation: the mean firing rates *r*_*PY R*_, *r*_*BiC*_, *r*_*CCK*_ and *r*_*PV*_ will be denoted by *x*_1_, *x*_2_, *x*_3_ and *x*_4_, respectively. The maximal firing rates *r*_0,*P Y R*_, *r*_0,*BiC*_, *r*_0,*CCK*_ and *r*_0,*P V*_ will be denoted by *r*_1_, *r*_2_, *r*_3_ and *r*_4_, respectively. To understand the mechanism of PRM we break it into three subsystems: PYR+BiC, PYR+PV and CCK+PV. The first considers the dynamics of PYR and BiC cells:

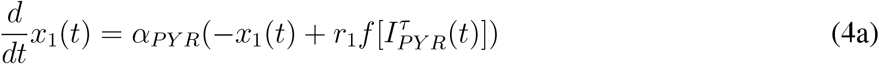

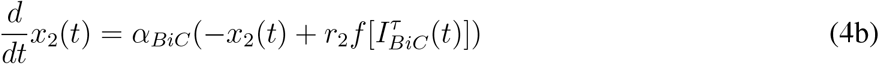

where 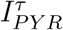 is as in Eq. (3a) with *w*_*PV* →*PY R*_ = 0 and 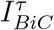 is as in Eq. (3b). The second one considers the dynamics of PYR and PV:

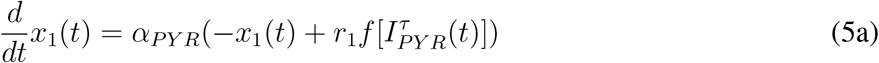

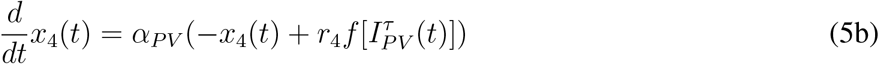

where 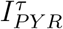 and 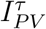 are defined as in Eq. (3a) and Eq. (3d), respectively, with *w*_*BiC*→*PY R*_ = 0 and *w*_*CCK*→*PV*_ = 0. The third subsystem considers the dynamics of PV and CCK, and is given by:

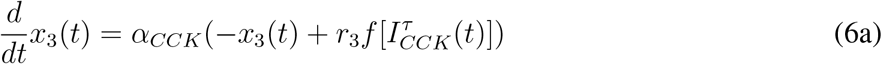

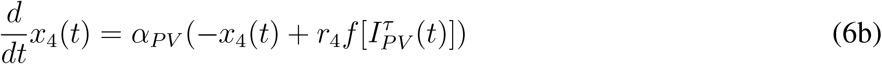

where 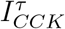 and 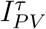 are defined as in Eq. (3c)-(3d), respectively, with *w*_*PY R*→*PV*_ = 0.

We start the analysis with the system of Eqs. (4a)-(4b). The point 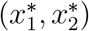 is an equilibrium point of Eqs. (4a)-(4b) if there is a solution to

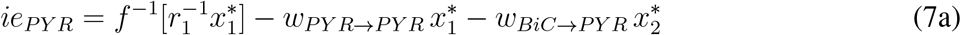

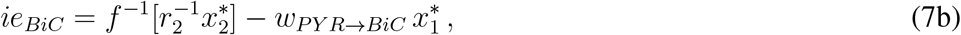

where *f*^−1^(*y*) = *β*^−1^ ln(*y/*(1 − *y*)). Note that we have dropped the additional inputs *STIM*_*PY R*_ and *STIM*_*BiC*_ from Eqs. (7a)-(7b) as they were initially considered to be zero.

Denote the right hand side of Eq. (4a) by *ϕ*(*x*_1_, *x*_2_), and of Eq. (4b) by *χ*(*x*_1_, *x*_2_). Linearizing Eqs. (4a)-(4b) around the equilibrium point 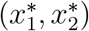 we find the Jacobian matrix

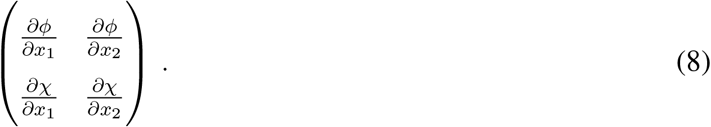

The first-order partial derivatives of the above matrix are

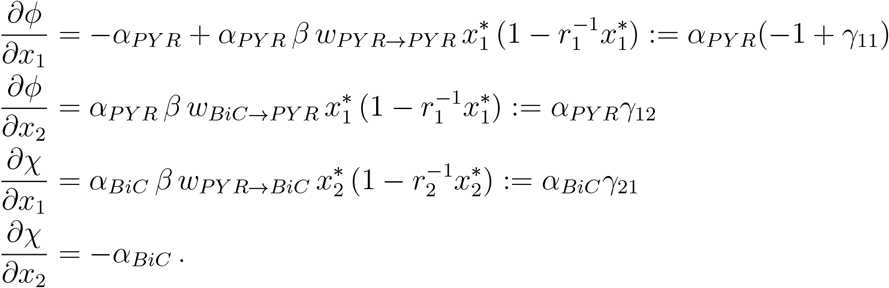

The solution of the linearized system is of the form (*x*_1_(*t*), *x*_2_(*t*)) = (*c*_1_, *c*_2_)*e*^*λt*^ where *c*_1_, *c*_2_ are nonzero constants, *λ* are the eigenvalues and *t >* 0. A condition on *λ* is given by the equation

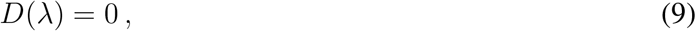

where

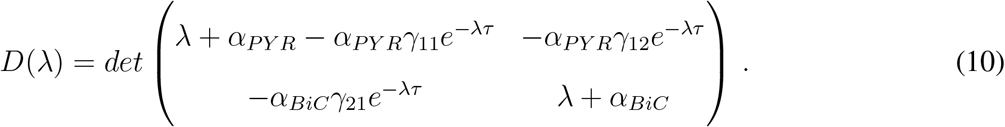

From Eq. (9) we have

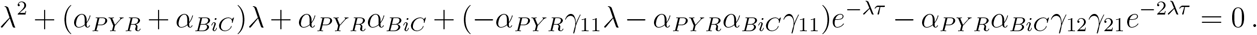

To simplify the above expression we multiply by *e*^*λτ*^ :

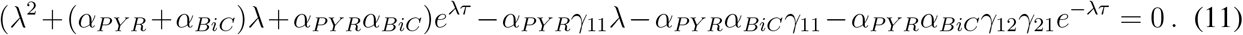

If *λ* = *iω*, for *ω* a nonzero real number, the bifurcation condition is defined by the simultaneous solution of the equations Re(*D*(*iω*)) = 0 and Im(*D*(*iω*)) = 0. Substituting *λ* = *iω* into Eq. (11) we conclude that the real and imaginary parts are:

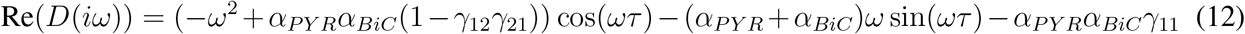

and

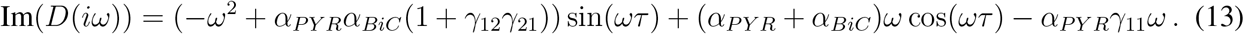

Taking Re(*D*(*iω*)) = 0 and Im(*D*(*iω*)) = 0 we find

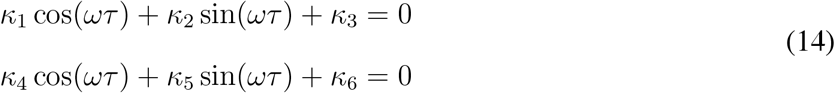

where we have further simplified the notation as follows:

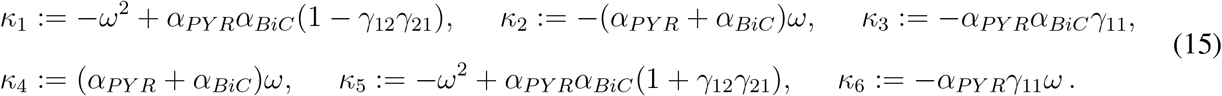

From Eq. (14) we have that

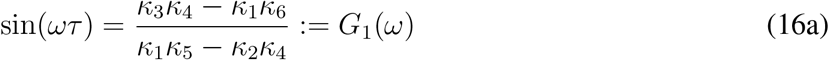

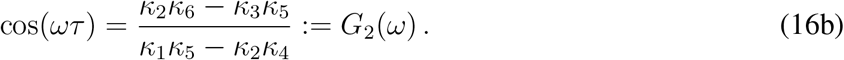

Notice that *G*_1_ and *G*_2_ depend only on *ω*, hence solving Eqs. (16a)-(16b) with respect to *ω* we are able to find the frequency. Indeed, the trigonometric equation sin^2^(*ωτ*) + cos^2^(*ωτ*) = 1 implies that 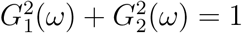. Solving the latter equation with parameter values of *Set 0* we found a frequency in the theta band. From Eq. (16a) we may also calculate the delay *τ* as follows:

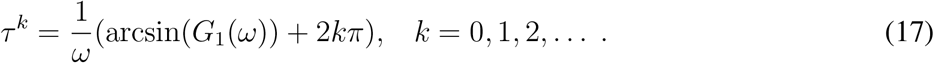

The smallest such value is the first bifurcation point.

We repeat the same steps as above for the stability analysis of the two other subsystems (PYR+PV and CCK+PV). In FIGURE 3G, we plot the bifurcation points (coloured triangles) for each of the three subsystems. For delays smaller than the critical ones the subsystems are either stable or oscillations are damping. For delays larger than the critical ones the oscillations are periodic. The curves plotted in FIGURE 3G are approximations of the frequencies at given delays, as computed by Eq. (17).

## SUPPORTING INFORMATION

There are Supplementary FIGURES and TABLES.

*SUPPL FIGURE 1* **Intrinsic firing rates for the four cell types in the PRM**.

*SUPPL FIGURE 2*: **Additional distributions**.

*SUPPL FIGURE 3*: **Example illustration of slope quantification for theta and gamma powers and theta frequency with STIM variation**.

*SUPPL FIGURE 4*: **Example illustration of PYR activities and their power spectral densities**.

*SUPPL FIGURE 5*: **Changes in regularity of theta events**.

*SUPPL FIGURE 6*: **A blow up of activities to illustrate timings**.

*SUPPL TABLE 1*: **Synaptic weight values and characteristics for the ten selected parameter sets**

*SUPPL TABLE 2*: **Theta and gamma power changes with connection removals**

*SUPPL TABLE 3*: **Changes in amplitude and interspike intervals of LFP theta with and without CCK**

*SUPPL TABLE 4*: **Statistical comparisons between cell types for theta and gamma power and ranges**

*SUPPL TABLE 5*: **Statistical comparisons between cell types for theta frequency**

