## Supplemental Figures and Tables for "Cell-type specific contributions to theta-gamma coupled rhythms in the hippocampus"

*Journal:* **NETWORK NEUROSCIENCE**

RESEARCH

### Supporting Information

### *SUPPLEMENTARY FIGURES (6) AND SUPPLEMENTARY TABLES (5)*

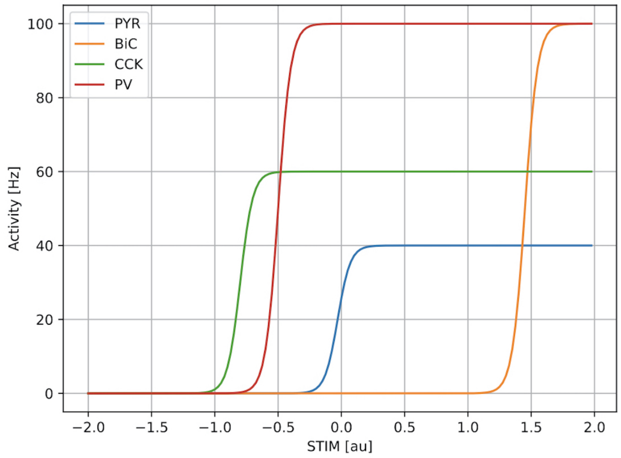


**Figure 1. Intrinsic firing rates for the four cell types in the PRM.**

As STIM is varied, the firing of the four cell types changes as shown.

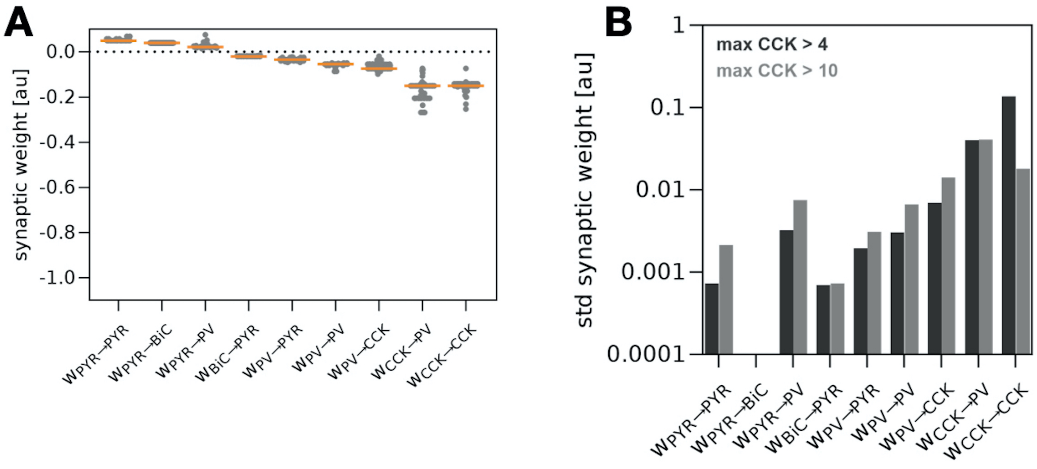


**Figure 2. Additional distributions.**

**A.** Distribution of synaptic weights ($w$) for each of the nine different connections in the PRM, from 200 constrained parameter sets of $w$ found when running the genetic algorithm for a maximal firing of CCK constrained to be greater than 10 (compare with FIGURE 2**A** in main text when maximal firing of CCK is 4). The orange line depicts the median. **B.** Bar graph showing the standard deviations (std) of $w$ for each of the nine different connections when maximal firing of CCK was either 4 or 10 as labelled. As expected, maximal CCK firing greater than 4 or 10 results in different $w$'s, but the relative sizes of $w$ remained the same, and the CCK→PV connection remained as a $w$ with one of the largest std.

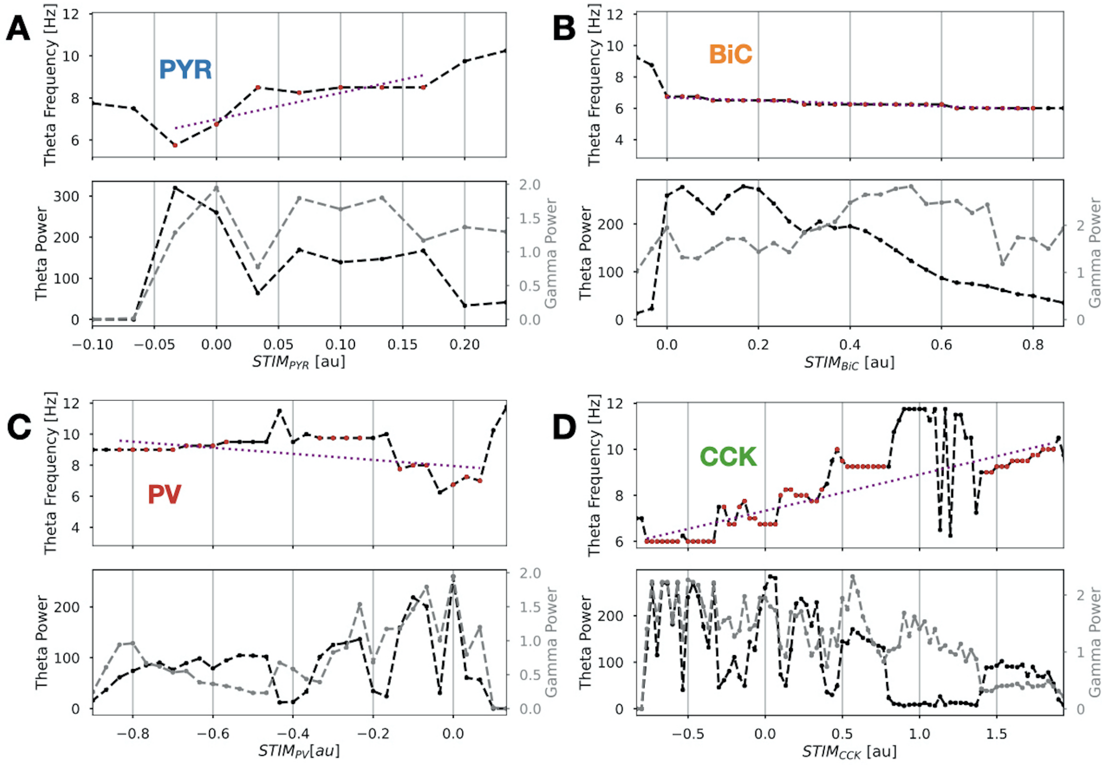


**Figure 3. Example illustration of slope quantification for theta and gamma powers and theta frequency with STIM variation.**

LFP theta frequency (top parts) and theta and gamma power (bottom parts) in the four-cell system with varying STIM to PYR (**A**), BiC (**B**), PV (**C**), and CCK (**D**) cell types are plotted using *Set 4* parameter values. The plots are truncated to be centred around the values of STIM that allow the system to exhibit theta-gamma coupled rhythms, and one can immediately see that there is a wider range (i.e., difference between the maximum and minimum) of STIM values for CCK relative to the other cell types. This is summarized in FIGURE 4**D** in main text. The STIM values where sufficient theta-gamma coupled rhythms occur are marked using red dots on the theta frequency plots. The dashed purple line shows the line best fitting the variation of theta frequency with STIM using only the points (red dots) where theta-gamma oscillations occur. Similar lines (not shown) and fits were done for theta and gamma powers. The slopes of the theta and gamma power line fits are used to make the plots shown in FIGURE 4**A, B** in main text. The plot using the slopes of the theta frequency line fits is shown in FIGURE 4**C** in main text. We note that changes in theta and gamma powers and theta frequency do not necessarily change in a smooth fashion. This jaggedness is due to the dynamic, nonlinear coupling interactions occurring between theta and gamma rhythms which affects the amplitude and expression of the theta-gamma coupled rhythms as STIM is varied, and hence theta and gamma powers and theta frequency. We illustrate this in SUPPLEMENTARY FIGURE 4.

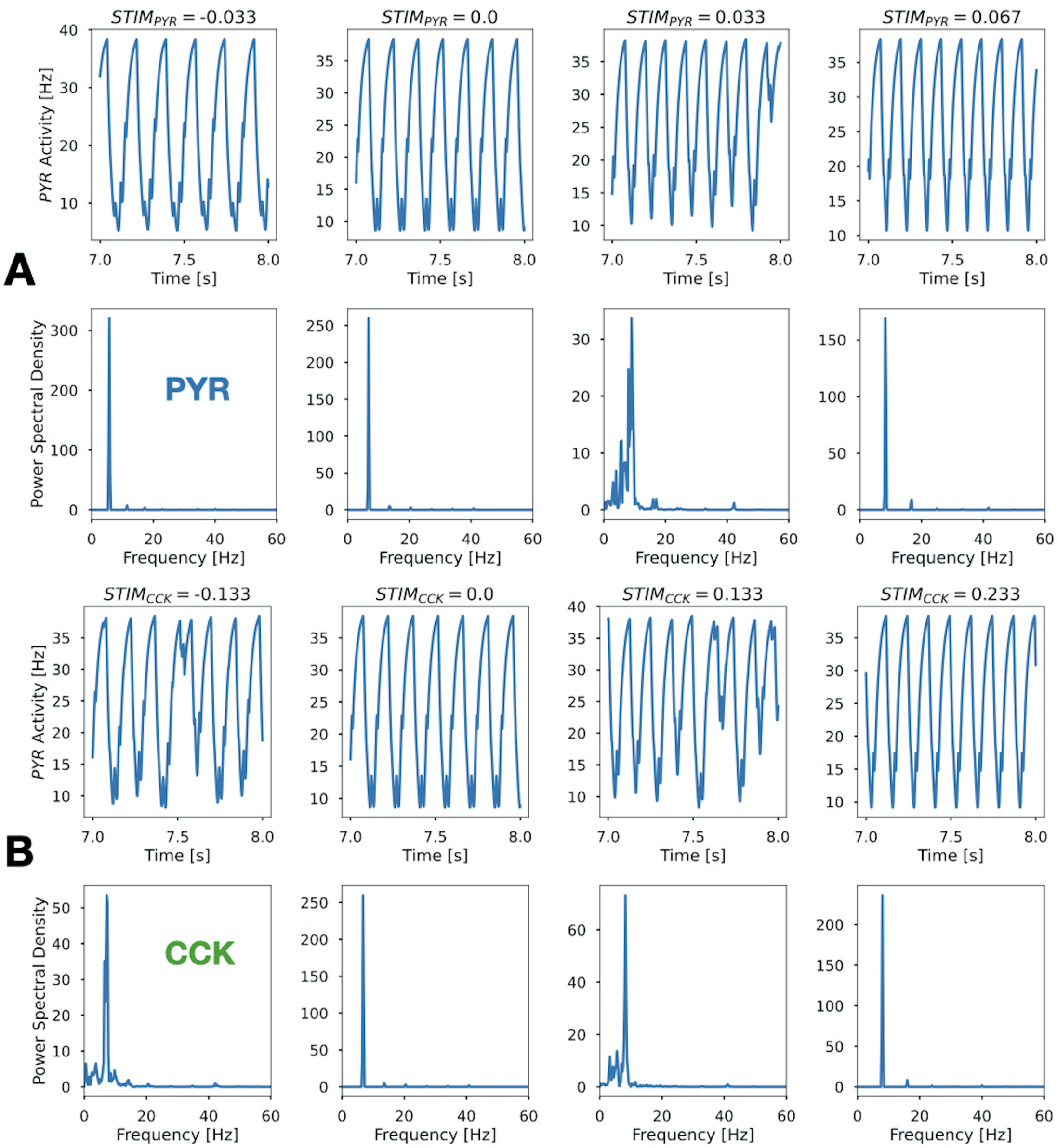


**Figure 4. Example illustration of PYR activities and their power spectral densities.**

PYR activity outputs for four different STIM values and the power spectral density (PSD) associated with the PYR activity (LFP representation). **A.** STIM values when applied to PYR **B.** STIM values when applied to CCK. This is for *Set 4* parameter values, representing specific red dots shown in SUPPLEMENTARY FIGURE 3. From these selected examples, it is clear why there is jaggedness in the theta and gamma powers and theta frequency. In viewing the PSDs, it is clear that the theta powers are much larger than the gamma powers. PSD units are Hz.

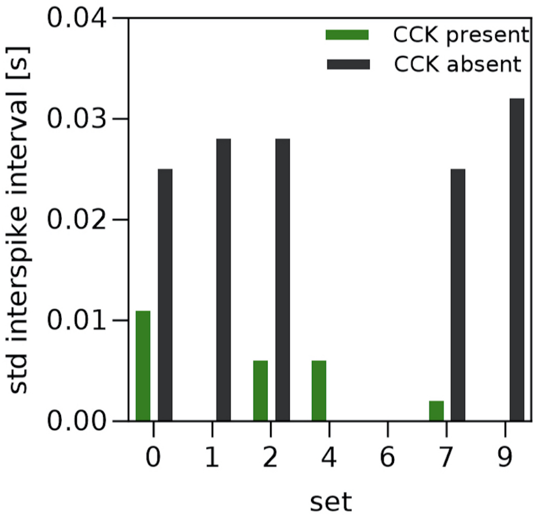


**Figure 5. Changes in regularity of theta events.**

Bar graph showing how the theta rhythm regularity is affected by the presence of absence of CCK during ongoing theta-gamma coupled rhythms for seven of the ten sets as labelled. If no bar is present, then the standard deviation (std) is zero. The specific values are given in SUPPLEMENTARY TABLE 3.

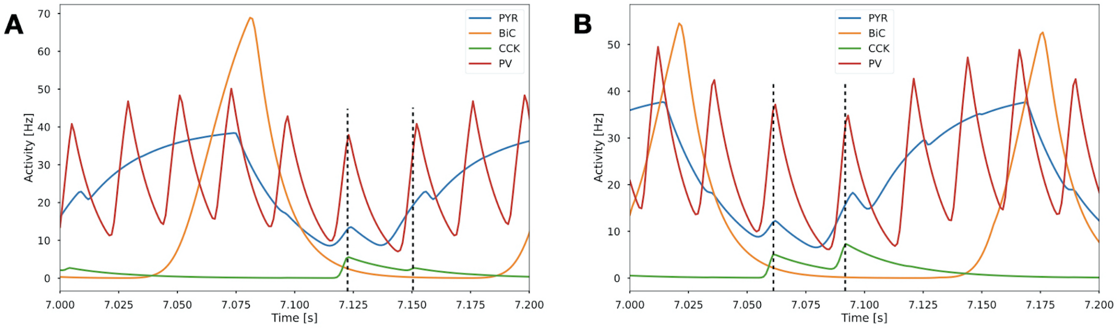


**Figure 6. A blow up of activities to illustrate timings.**

Activities of the four cell types are shown for a 0.1 second range for *Set 4* parameter values (**A**), and *Set 5* parameter values (**B**). Vertical lines show that the peaks of CCK precede both PV and PYR, thus illustrating support for the two-phase process schematized in FIGURE 6**C** (in main text) where CCK initiates the gamma in the first phase.

**Table 1. Synaptic weight values and characteristics for the ten selected parameter sets**

| **Set** | $w_{PYR \to PYR/BiC/PV}$ *(au)* | $w_{BiC \to PYR}$ *(au)* | $w_{CCK \to CCK/PV}$ *(au)* | $w_{PV \to PV/PYR/CCK}$ *(au)* | *PV/BiC ratio* | *max CCK (Hz)* | *Theta freq (Hz)* |
|---|---|---|---|---|---|---|---|
| 0 | 0.05/0.04/0.02 | -0.02 | -0.15/-0.15 | -0.055/-0.03/-0.075 | 0.71 | 10.11 | 7.75 |
| 1 | 0.05/0.04/0.02 | -0.02 | -0.15/-0.107 | -0.055/-0.03/-0.075 | 0.71 | 6.29 | 6.5 |
| 2 | 0.05/0.04/0.02 | -0.019 | -0.15/-0.15 | -0.055/-0.03/-0.075 | 0.71 | 8.08 | 8.0 |
| 3 | 0.05/0.04/0.036 | -0.02 | -0.15/-0.107 | -0.055/-0.03/-0.075 | 1.05 | 7.38 | 4.5 |
| 4 | 0.05/0.04/0.02 | -0.019 | -0.15/-0.078 | -0.055/-0.03/-0.075 | 0.72 | 5.71 | 6.75 |
| 5 | 0.05/0.04/0.02 | -0.02 | -0.15/-0.11 | -0.055/-0.035/-0.075 | 0.83 | 8.18 | 6.5 |
| 6 | 0.05/0.04/0.02 | -0.02 | -0.15/-0.024 | -0.055/-0.025/-0.075 | 0.71 | 4.76 | 8.5 |
| 7 | 0.05/0.04/0.02 | -0.02 | -0.48/-0.078 | -0.055/-0.03/-0.075 | 0.73 | 5.45 | 6.75 |
| 8 | 0.05/0.04/0.027 | -0.019 | -0.27/-0.078 | -0.055/-0.03/-0.052 | 0.85 | 9.92 | 7.0 |
| 9 | 0.05/0.04/0.02 | -0.019 | -1.03/-0.086 | -0.055/-0.03/-0.071 | 0.72 | 5.46 | 6.75 |

*All of the synaptic weight values for the ten parameter sets are given. Other parameter values are given in TABLE 1 (in main text). Unless otherwise indicated, STIM is set to zero for all cell types. Also shown are the activity ratios of PV/BiC and the maximal CCK activities which formed part of the constraints (see details in Model and methods section), as well as the theta frequency, for each of the ten parameter sets.*

**Table 2. Theta and gamma power changes with connection removals**

| **Set** | *Connection type removal* | *Theta power ratio*[1] | *Gamma power ratio*[1] | *Sufficient?*[2] *(Y/N)* |
|---|---|---|---|---|
| 0-3,5 | PYR→PYR* | $\approx 0$ | $> 0.2$ | N |
| 4 | PYR→PYR* | $\approx 0$ | 0.06 | N |
| 6-9 | PYR→PYR* | $\approx 0$ | $< 0.05$ ($\neq 0$) | N |
| 0-9 | PYR→BiC | $\approx 0$ | $\approx 0$ | N |
| 0,2 | PYR→PV | $\approx 0$ | $\approx 0$ | N |
| 1,3 | PYR→PV | $> 0.2$ | $> 0.2$ | N |
| 4-9 | PYR→PV | $> 0.2$ | $> 0.2$ | **Y** |
| 0-9 | BiC→PYR* | $\approx 0$ | $\approx 0$ | N |
| 0-9 | PV→PYR* | $\approx 0$ | $\approx 0$ | N |
| 0-9 | PV→PV | $\approx 0$ | $\approx 0$ | N |
| 0,2,3,5 | PV→CCK | $< 0.07$ ($\neq 0$) | $> 0.2$ | N |
| 1 | PV→CCK | 0.11 | $> 0.2$ | N |
| 4 | PV→CCK | 0.17 | $> 0.2$ | N |
| 6,9 | PV→CCK | $> 0.2$ | $> 0.2$ | **Y** |
| 7 | PV→CCK | $> 0.2$ | $> 0.2$ | N |
| 8 | PV→CCK | 0.12 | $> 0.2$ | N |
| 0-9 | CCK→PV* | $\approx 0$ | $< 0.01$ ($\neq 0$) | N |
| 0-5,7-9 | CCK→CCK | $\approx 0$ | $\approx 0$ | N |
| 6 | CCK→CCK | $> 0.2$ | $> 0.2$ | **Y** |

[1] *Theta or gamma power ratio is the ratio of the theta power when the particular connection is removed divided by the reference theta or gamma power when no connections are removed.*

[2] *Sufficient means that both theta and gamma powers are large enough to be considered to have theta-gamma coupled rhythms to be present.* $> 25\%$ *of the reference power was used as the threshold. Note that if* $> 20\%$ *was used, more cases would have been considered to be sufficient.*

**Due to applied constraints (see FIGURE 1* ***E-H*** *in main text), removal of these connections is expected to yield no theta rhythms, and hence no theta-gamma coupled rhythms. See Model and methods section for further details.*

**Table 3. Changes in amplitude and interspike intervals of LFP theta with and without CCK**

| **Set** | *Amplitude (au)* | | *Mean ISI (s)* | | *Std ISI (s)* | |
|---|---|---|---|---|---|---|
| | *CCK present* | *CCK absent* | *CCK present* | *CCK absent* | *CCK present* | *CCK absent* |
| 0 | 38.2 | 38.1 | 0.128 | 0.147 | 0.011 | 0.025 |
| 1 | 38.5 | 37.9 | 0.153 | 0.135 | 0.0 | 0.028 |
| 2 | 38.2 | 38.1 | 0.126 | 0.146 | 0.006 | 0.028 |
| 4 | 38.4 | 38.4 | 0.146 | 0.167 | 0.006 | 0.0 |
| 6 | 38.3 | 38.3 | 0.117 | 0.117 | 0.0 | 0.0 |
| 7 | 38.2 | 37.9 | 0.147 | 0.140 | 0.002 | 0.025 |
| 9 | 38.4 | 38.3 | 0.147 | 0.167 | 0.0 | 0.032 |

*ISI = interspike interval between PYR peaks (i.e., theta cycles of LFP representation); Std = standard deviation.*

**Table 4. Statistical comparisons between cell types for theta and gamma power and ranges**

| ***Theta power*** | | | ***Gamma power*** | | | **STIM range** | | |
|---|---|---|---|---|---|---|---|---|
| *Cell 1* | *Cell 2* | *p-value* | *Cell 1* | *Cell 2* | *p-value* | *Cell 1* | *Cell 2* | *p-value* |
| PYR | BiC | $6.3 \times 10^{-2}$ * | PYR | BiC | $1.9 \times 10^{-2}$ | CCK | PYR | $1.1 \times 10^{-5}$ |
| PYR | CCK | $1.1 \times 10^{-5}$ | PYR | CCK | $1.5 \times 10^{-3}$ | CCK | BiC | $2.2 \times 10^{-5}$ |
| PYR | PV | $1.1 \times 10^{-5}$ | PYR | PV | $4.9 \times 10^{-4}$ | CCK | PV | $2.2 \times 10^{-5}$ |
| BiC | PV | $1.1 \times 10^{-5}$ | PV | BiC | $1.9 \times 10^{-1}$ * | | | |
| BiC | CCK | $1.1 \times 10^{-5}$ | PV | CCK | $1.9 \times 10^{-2}$ | | | |
| PV | CCK | $1.1 \times 10^{-5}$ | BiC | CCK | $6.8 \times 10^{-1}$ * | | | |

** Not statistically significant. These comparisons are related to FIGURE 4**A,B,D** in main text. Comparisons were done using Mann-Whitney (95%, p<0.05).*

**Table 5. Statistical comparisons between cell types for theta frequency**

| | ***Theta frequency*** | |
|---|---|---|
| *Cell 1* | *Cell 2* | *p-value* |
| PYR | BiC | $3.3 \times 10^{-4}$ * |
| PYR | CCK | $2.1 \times 10^{-3}$ |
| PYR | PV | $1.1 \times 10^{-5}$ |
| BiC | CCK | $7.5 \times 10^{-2}$ * |
| BiC | PV | $3.9 \times 10^{-3}$ |
| CCK | PV | $1.1 \times 10^{-5}$ |

** Not statistically significant. These comparisons are related to FIGURE 4**C** in main text. Comparisons were done using Mann-Whitney (95%, p<0.05).*
